# textToKnowledgeGraph: Generation of Molecular Interaction Knowledge Graphs Using Large Language Models for Exploration in Cytoscape

**DOI:** 10.1101/2025.07.17.664328

**Authors:** Favour James, Christopher Churas, Dexter Pratt, Augustin Luna

**Affiliations:** Department of Electronic and Electrical Engineering, Obafemi Awolowo University; Department of Medicine, University of California, San Diego, La Jolla, CA, United States; Computational Biology Branch, National Library of Medicine and Developmental Therapeutics Branch, National Cancer Institute

**Keywords:** Large Language Models, Retrieval Augmented Generation, Knowledge Graphs, Biological Networks

## Abstract

**Motivation:** Knowledge graphs (KGs) are powerful tools for structuring and analyzing biological information due to their ability to represent data and improve queries across heterogeneous datasets. However, constructing KGs from unstructured literature remains challenging due to the cost and expertise required for manual curation. Prior works have explored text-mining techniques to automate this process, but have limitations that impact their ability to capture complex relationships fully. Traditional text-mining methods struggle with understanding context across sentences. Additionally, these methods lack expert-level background knowledge, making it difficult to infer relationships that require awareness of concepts indirectly described in the text. Large

Language Models (LLMs) present an opportunity to overcome these challenges. LLMs are trained on diverse literature, equipping them with contextual knowledge that enables more accurate extraction of information.

**Results:** We present textToKnowledgeGraph, an artificial intelligence tool using LLMs to extract interactions from individual publications directly in Biological Expression Language (BEL). BEL was chosen for its compact and detailed representation of biological relationships, allowing for structured and computationally accessible encoding. This work makes several contributions. 1. Development of the open-source Python textToKnowledgeGraph package (pypi.org/project/texttoknowledgegraph) for BEL extraction from scientific articles, usable from the command line and within other projects, 2. An interactive application within Cytoscape Web to simplify extraction and exploration, 3. A dataset of extractions that have been both computationally and manually reviewed to support future fine-tuning efforts.

**Availability:** https://github.com/ndexbio/llm-text-to-knowledge-graph

## 1. Introduction

Artificial Intelligence (AI) chatbots and other applications frequently use external information stores to enable LLMs to answer questions requiring specific, reliable knowledge. One of the most common approaches is the RAG (Retrieval-Augmented Generation). RAG is a technique that enhances Large Language Models (LLMs) by dynamically retrieving relevant information from external knowledge sources and incorporating it into the model’s context during generation, thereby enabling more accurate and grounded responses (Wang, Zhou, et al., 2024). While RAG effectively retrieves relevant information, it still faces the constraint of the LLM’s context window size. When multiple documents or passages are highly relevant to a query, RAG must either selectively omit potentially important information or implement complex chunking and summarizing strategies, which can impact the completeness and accuracy of responses. Recently, there has been a growing interest in using knowledge graphs as a source of information for AI systems. A knowledge graph (KG) is a structured network that represents relationships between concepts, entities, or pieces of information, typically using nodes (for entities) and edges (for relationships) (Slater, 2014). KGs are well-known in fields like semantic web research and knowledge representation. Large general-purpose biomedical knowledge graphs are publicly available (Morris et al., 2023), but tools to enable biomedical researchers to construct topic-focused knowledge graphs remain limited. However, LLMs have recently grown increasingly capable of reliably extracting complex knowledge from documents, presenting an opportunity to make the construction of custom knowledge graphs dramatically easier.

By converting these unstructured texts into a structured KG, we can help LLMs focus on the specific information of scientific interest, enhancing both the precision and depth of their analyses (Liu et al., 2019; Feng et al., 2025). Such graphs must integrate various data types while maintaining clarity in their relationships and interactions. We present textToKnowledgeGraph, a Python package to extract complex, detailed knowledge of scientific interactions as graphs in the BEL (Biological Expression Language). We chose BEL because it can capture qualitative causal and correlative relationships between biological entities and processes in a structured and computationally accessible manner (Charles Tapley Hoyt et al., 2017). BEL can encode causal statements in a format that is amenable to both human and machine interpretation. Additionally, BEL uses less text for interaction representation than comparable formats, such as the Biological Pathway Exchange (BioPAX) format. This reduction in text reduces associated costs and may also minimize errors by simplifying task complexity (Slater, 2014; Wang, Zhou, et al., 2024).

Another reason for choosing BEL is its amenability to visualization; the visualization of networks can aid in organizing biological knowledge (Novère et al., 2009). Our textToKnowledgeGraph tool integrates directly with the widely used Cytoscape ecosystem (Charles Tapley Hoyt et al., 2017). By generating highly structured BEL output that flows seamlessly into Cytoscape as a network graph, textToKnowledgeGraph addresses the need for interoperable and analyzable knowledge graphs in biology. The Cytoscape ecosystem, comprising Cytoscape (Shannon et al., 2003), Cytoscape Web (Ono et al., 2025) for interactive network exploration, and Network Data Exchange (NDEx) for storing and sharing networks (Pratt et al., 2015), enables researchers to explore complex biological relationships. Our tool uses the CX2 JSON-based format, the default network exchange format for transferring content between Cytoscape core components.

## 2. Methods

### 2.1. What is Biological Expression Language (BEL)?

BEL follows a subject-predicate-object structure, where the subject and object represent biological entities, and the predicate indicates the type of interaction (Charles Tapley Hoyt et al., 2017). BEL represents complex entities such as the activity of a protein or a protein with post-translational modifications in expressions using functional composition. Entities are specified as expressions grounded by established life science identifiers in vocabularies, such as HGNC for genes and CHEBI for chemicals. Multiple BEL statements can be linked to form larger networks.

Example of a BEL Statement (Figure 1):

**Fig 1.**
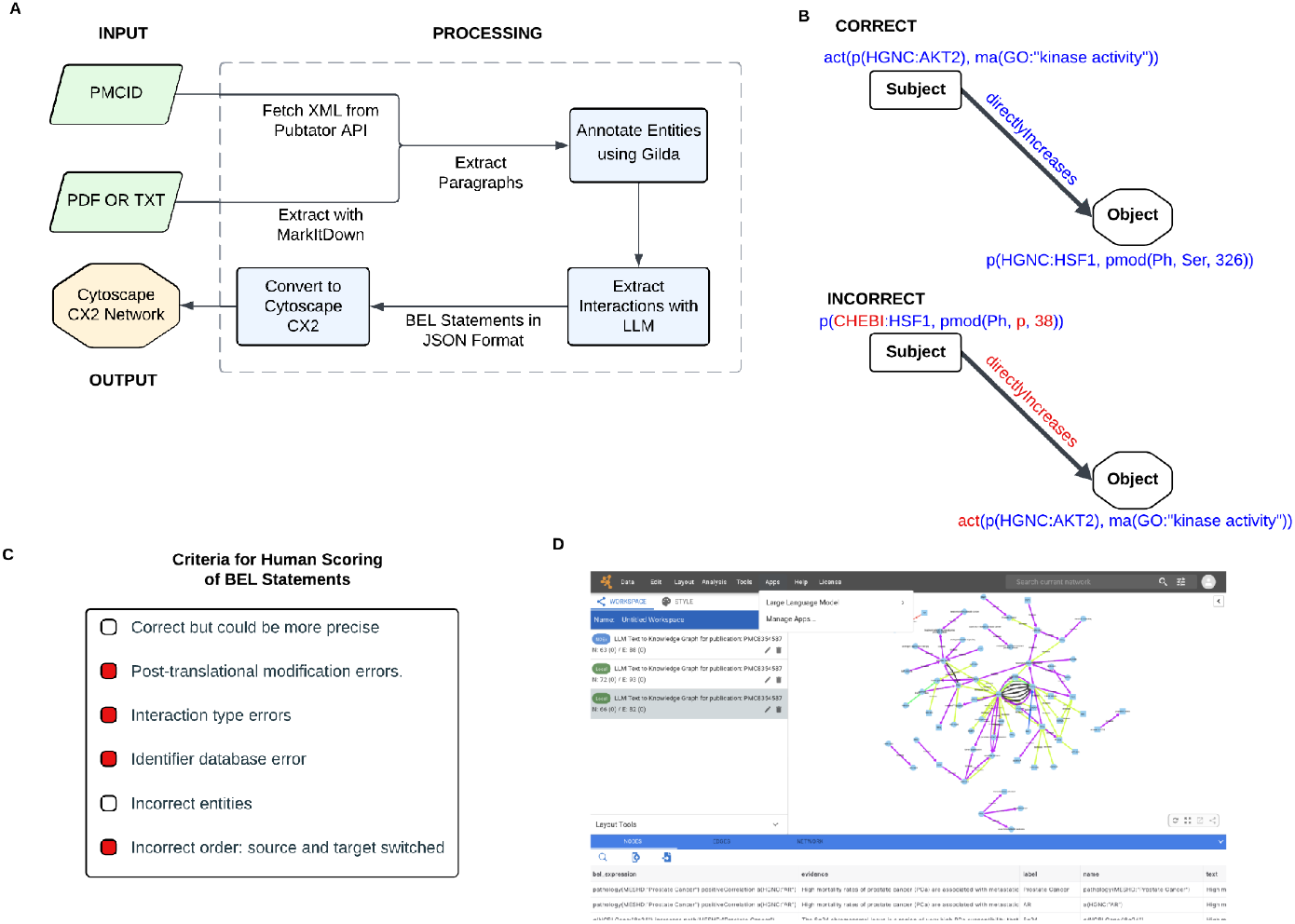
A) textToKnowledgeGraph processing pipeline, B) 2 representations of the same interaction (correct and incorrect with various error types), C) criteria used to evaluate the BEL statements; red boxes indicate the incorrect issues in sub-figures B and D) screenshot of our app in Cytoscape Web.

act(p(HGNC:AKT1),ma(GO:”kinaseactivity”)) directlyIncreases p(HGNC:HSF1, pmod(Ph, Ser, 326))

This translates to:

*The kinase activity of Akt1 protein directly causes an increase in the amount of HSF1 protein that is phosphorylated at SER 32*

### 2.2. Input and Output

#### Input (Figure 1A)

The primary input is either a PubMed Central (PMC) ID or a local file path. In the case of a PMCID, a pre-processed manuscript is obtained from PubTator, which provides entity and relation annotations for PubMed abstracts and full-text articles from the PMC open access subset (Wei et al., 2024). Alternatively, the Microsoft MarkItDown tool supports text extraction for various file formats. Credentials for the OpenAI API are also needed. Optional input includes NDEx credentials, allowing users to upload the CX2 output to NDEx.

#### Output (Figure 1A)

The pipeline produces two outputs:

1. <u>BEL as JSON:</u> BEL statements are stored in JavaScript Object Notation (JSON) format. We also include: the original paragraph from which the interactions were extracted, the BEL statement, and an evidence field containing the sentence within the paragraph from which the interaction was extracted.
2. <u>CX2 Format:</u> A format compatible with Cytoscape, NDEx, and other Cytoscape ecosystem tools.

### 2.3. Implementation

A major component of the textToKnowledgeGraph is the “prompt,” which guides the LLM in extracting and structuring interactions. The prompt provides the LLM with essential knowledge for creating BEL. The prompt (Available in the code repository) includes information on BEL syntax with examples and allowable identifiers.

For PMCID inputs, the tool retrieves the full article text in (Extensible Markup Language) XML format from PubTator (Wei et al., 2024), which is already broken down into paragraphs. For file uploads, the document is processed with an LLM to isolate relevant paragraphs while removing sections such as references and acknowledgments. The paragraphs from both sources are grounded using the Gilda service (Gyori et al., 2022) for standardized named entity recognition (NER) annotations. These paragraphs, along with their list of pre-annotated entities, are passed to the LLM to minimize LLM hallucinations regarding entities (e.g., avoiding the invention of non-standard identifiers). We leverage LangChain to create a reusable extraction chain that combines the prompt, the GPT-4o model, and an output parser. This chain is invoked on each paragraph, generating BEL statements. We validate our LLM output structure using Pydantic (pydantic.dev). Next, textToKnowledgeGraph converts the results into CX2 for visualization using the ndex2 package, which connects Cytoscape to NDEx. The extracted data is first structured using Pandas (The Pandas Development Team, 2020), then transformed into CX2 format via the PandasDataFrameToCX2NetworkFactory ndex2 Python package function, where each entity is assigned a unique node, and relationships are mapped with attributes. Installation and usage instructions can be found in the project repository.

## 3. Results

### 3.1. Evaluation of BEL Statements

We provide the results of our evaluations in the code repository as Jupyter notebooks. It is possible to make decisions on the quality of generated BEL statements (Fig. 1B). To evaluate the quality, 67 statements, which contained 75 nodes and 50 edges, were reviewed by 5 different reviewers following the error criteria (Fig. 1C). These reviews showed that identifier errors were often due to Gilda entity recognition. 62.8% of individual reviews were marked as correct and ‘correct but could be more precise’. In addition, 40% of statements had strong agreement, with at least four out of five reviewers judging them positively; this criterion was chosen because, in certain cases, there could be alternative representations for particular interactions.

We also compared our results with those of an established interaction extraction system, INDRA, by parsing 20 sentences with both tools and analyzing the overlap in results (Bachman et al., 2023). We found that textToKnowledgeGraph extracted more statements than INDRA from the sentences and that the tools provided many complementary rather than redundant outputs. Interactions extracted by INDRA were converted to BEL using pyBEL. Whereas our pipeline extracted 67 statements, INDRA produced only 27 statements for the same set of 38 unique evidence texts. An LLM-as-a-judge approach was applied to rate the similarity of statements extracted from the same text into three categories: Good, representing the same relationship; Medium, where statements are related but differ slightly in specificity; and Bad, where statements describe different relationships.

To evaluate the results between LLM and INDRA, we developed a scoring logic that compared statement components (e.g., subject, object) from INDRA with those from our tool. Statements with no matching components were given a score of 0. Components were given (e.g., if the relationship matched a sub-score of 0.4 was given); the highest score possible was 1. Statements with the highest score were considered the best match to the LLM. The statements from INDRA and our tool were then compared to assess their similarity using the LLM-as-judge, which evaluates semantic similarity due to the possibility of alternative valid representations. We identified 29 evidence texts where both textToKnowledgeGraph and INDRA produced BEL statements. In a component-level analysis of these pairs, 6 out of 29 pairs (20.7%) had matching subjects, 11 (37.9%) had matching relationship types, and 8 (27.6%) had matching objects. The namespace agreement was higher, with 16 pairs (55.2%) matching for the subject namespace and 15 pairs (51.7%) for the object namespace. Overall, 8 out of 29 statements (27.6%) were rated as ‘Good’ matches by the LLM-as-judge scoring.

### 3.2. Use Cases

#### 3.2.1. Cytoscape Web Application

Cytoscape Web supports textToKnowledgeGraph to simplify the use of our tool. Cytoscape users can input a PMCID to process it using textToKnowledgeGraph and can save it to NDEx (screenshot in Fig. 1D) from within the app.

#### 3.2.2. Graph Retrieval Augmented Generation (GraphRAG)

As an example, we demonstrate how KGs generated by our tool can be utilized for retrieval-augmented generation purposes using a GraphRAG approach, in which a query to our KG provides additional relevant information to be incorporated into the prompt context. We provide a GraphRAG pipeline as a Jupyter notebook that retrieves the 1-hop neighborhood of specified genes from a BEL KG stored in NDEx, and injects those BEL statements into the LLM prompt in CX2 JSON representation. The LLM then generates a text response containing biological relationships represented in the KG. For demonstration, we show that the answer to “How does metabolism affect DNA damage response?” is modified and clarified as KGs are introduced. This use case was chosen as it allows us to showcase how KGs can be merged to describe connections between different biological processes.

More specifically, the notebook shows seven variations of posing the query “How does metabolism affect DNA damage response?” to an LLM. We conduct demonstrations for two different knowledge graphs (one derived from a paper focused on DNA damage and another related to metabolism), as well as a merged scenario using both knowledge graphs, which drives the LLM response. Each query was performed with and without asking the LLM to utilize its internal knowledge, and one final case was performed without any KG. This enables the model to utilize both the KG and its internal knowledge to enrich and focus responses (e.g., our example specifically focuses on the NAD^+^–SIRT1–PARP1 axis that bridges DNA damage and metabolism), with text traceable to BEL statements in the merged network.

## 4. Conclusion

We develop textToKnowledgeGraph, which is accessible programmatically and via Cytoscape Web, that allows users to generate knowledge graphs from scientific literature using BEL. We evaluate this work through manual review and in comparison to similar tools (i.e., INDRA). Our results demonstrate that LLMs can extract more biological interactions than traditional pipelines. A key limitation observed during human review was that accurate entity recognition remains a crucial factor: misannotations from the Gilda service sometimes led to incorrect gene names, which directly impacted the quality of the extracted interactions. This highlights that while prompt design and LLM output structure are crucial, the reliability of upstream named entity recognition remains vital and needs continued improvement. Lastly, although it is outside the scope of the current effort, our approach is adaptable to other formats that capture biological interactions, and this work is a promising topic for investigation in future studies. Overall, our study suggests that combining LLM contextual capabilities with robust entity recognition and precise evaluation can advance automated knowledge graph construction.

## Funding

Financial support was provided by the National Institutes of Health (NIH) under grants R01HG009979, U19AI135990, U54CA209891, and U24CA184427. The National Resource for Network Biology (NRNB) from the National Institute of General Medical Sciences (NIGMS P41GM103504), and supported in part by the Division of Intramural Research (DIR) of the National Library of Medicine (NLM) (ZIALM240126), and Google Summer of Code.

## Acknowledgments

We thank the Cytoscape development team members, Jing Chen, Dylan Fong, and Chengzhan Gao, for their help with Cytoscape Web integration and internal tools for BEL review. We thank Ben Gyori for his advice on using INDRA. We also thank Clara Hu, Nicole Mattson, JungHo Kong, Patrick Wall, and Zhao Xiaoyu for their help with manual reviews of the generated BEL statements.

## Notes

https://github.com/ndexbio/llm-text-to-knowledge-graph

### Competing Interest Statement

The authors have declared no competing interest.

